# Endothelial Cell-Specific MCPIP Deletion Causes Endothelial Dysfunction and Impairs Post-ischemic Angiogenesis in Vivo

**DOI:** 10.1101/208884

**Authors:** Jianli Niu, Nidhi Kapoor, Jian Liang, Zhuqing Jin, Edilu Becerra, Pappachan E. Kolattukudy

## Abstract

Vascular endothelial cells (ECs) play an important role in angiogenesis and inflammatory responses. MCPIP (also known as Zc3h12a or Regnase-1), a newly identified suppressor of cytokine signaling, is expressed in endothelial cells. To directly test the role of endothelium-derived MCPIP in cardiovascular pathophysiology, we specifically targeted deletion of the murine MCPIP gene in the endothelium by using the loxP/Cre system. A floxed MCPIP knock in mouse line was crossbred with VEcadherin5-Cre mice to generate offspring with deletion of the MCPIP gene in ECs (ECKO). Ablation of MCPIP in ECs resulted in systemic inflammation, anemia, splenomegaly, increased vessel permeability, muscle wasting, endothelial dysfunction, thrombus formation, impaired blood perfusion, and reduced lifespan in these mice. Serum levels of cytokines, chemokines, and biomarkers of EC dysfunction were significantly elevated in the ECKO mice, suggesting a hypercytokinemia. These mice also were more susceptible to lipopolysaccharide-induced death. When subjected to ischemia, these mice showed defective post-ischemic angiogenesis and impaired blood flow recovery in hind limb ischemia and stroke models, as well as in wound healing. This effect was associated with an increased cellular infiltration, cytokine expression, and decreased angiogenic factors. Moreover, MCPIP–deficient ECs displayed decreased vascular sprouting and tube elongation in ex vivo aortic ring assay. MCPIP-knockdown in cultured ECs enhanced NF-κΒ activity and dysregulated synthesis of microRNAs linked with elevated cytokines and biomarkers of EC dysfunction. These data show, for the first time, that constitutive expression of MCPIP in ECs is essential to maintain ECs in a quiescent state by serving as an important negative feedback regulator that keeps the inflammatory signaling suppressed.

## Introduction

Vascular endothelium is recognized as a key player in the host’s immune responses to different physical and chemical stimuli.^1^ A critical initiating event in this process is endothelial cell (EC) activation, which is an immunological and inflammatory response that causes endothelial dysfunction characterized by increased expression of adhesion molecules, secretion of cytokines, change in phenotype from antithrombotic to prothrombotic, elicitation of T-cell-mediated immune response, and loss of vascular integrity.^2^ It is well established that endothelial dysfunction is a major determinant of the initiation and/or progression of many cardiovascular diseases, including atherosclerosis, congestive heart failure, diabetes mellitus and other inflammatory syndromes.^2^ EC activation/dysfunction is typically driven by transcription factor nuclear factor κB (NF-κΒ), which not only plays a key role in the induction of prothrombotic and pro-inflammatory responses, but also renders the endothelium more susceptible to apoptosis.^3-5^ Emerging evidence has also revealed that noncoding microRNAs constitute a new class of intra- and intercellular signaling molecules to modulate inflammation in ECs. ^6-8^

MCP-1-induced protein (MCPIP, also known as *Zc3h12a* or *Regnase*-1), originally discovered as a novel zinc-finger protein induced by monocye chemoattractant protein-1(MCP-1), was identified as a negative feedback inhibitor of cytokine signaling.^9,10^ So far, MCPIP has been implicated in the regulation of levels of cytokines, including interleukin (IL)-6, IL-12, IL-1β and IL-17. MCPIP not only functions as a suppressor of NF-κB signaling, ^11^ but has also been shown to disrupt other inflammatory pathways by regulating mRNA degradation,^12^ microRNA synthesis^13^ and IL-17 receptor degradation.^14^ Most notably, although MCPIP-deficient mice die around 8-10 weeks postnatally due to systemic inflammatory responses, ^12^ mice with deficiency of MCPIP in myeloid cells develop normally and have no overt phenotype.^15^ Expression of MCPIP has been observed in murine and human atherosclerotic lesions and rheumatoid arthritis.^16, 17^ Although we previously showed MCPIP regulates expression of vascular cell adhesion molecule 1 (VCAM)-1 and angiogenesis in cultured ECs, ^16,18, 19^ the pathophysiological role of MCPIP in the cardiovascular system remains largely unknown. In the present study, we investigated the role of MCPIP in the cardiovascular system *in vivo* based on analysis of mice with MCPIP deletion specifically in murine vascular ECs. Data presented here demonstrate that constitutive expression of MCPIP is essential to keep ECs in a quiescent state by suppressing NF-κB activation and microRNA synthesis.

## Materials and Methods

All animal procedures and protocols used in this study were approved by the University of Central Florida Animal Care and Use Committee, in accordance with the Guide for the Care and Use of Laboratory Animals published by the US National Institutes of Health (NIH Publication No.85-23, revised 2011). The MCPIP gene in ECs was disrupted by mating MCPIP^LoxP/LoxP^ mice, ^15^ with mice carrying the Cre recombinase gene driven by the EC-specific VEcadherin5 promoter (Jackson Laboratory, Stock #017968). A detailed, expanded Methods section is provided in the Online Supplemental Material that includes descriptions of the multiple analytic profiling, vascular permeability analyses, ex vivo mouse aortic ring assay, mouse middle cerebral artery occlusion model, mouse femoral artery ligation model, full-thickness punch wound model, laser Doppler perfusion imaging, cell culture, histological and histomorphometric assessments, fluorescent immunohistochemical analyses, cell culture, siRNA interfering, NF-κΒ luciferase reporter assay, quantitative real-time RT-PCR (qRT-PCR) and immunoblotting, and statistical analyses.

## Results

### Generation and phenotypic characterization of mice with EC-specific deletion of MCPIP

To ablate MCPIP expression in ECs, homozygous floxed MCPIP mice^15^ were crossed with the VEcadherin5-Cre mouse line (Jackson Laboratory, Stock #017968), and the progeny were bred to produce mice which are homozygous for endothelial-specific knockout of MCPIP (ECKO). ECKO mice on a C57BL/6 background were phenotypically normal at birth and nearly indistinguishable during the first three weeks of life. Thereafter, their growth was slower than their littermate controls. The adult ECKO animals were significantly smaller compared to age-matched wild type littermates. They survive for about 10-14 weeks. To gain insight into the causes of these phenotypic differences, sex- and age-matched wild type and ECKO mice were used for gross necropsies to determine the size and weight of major organs. Comparison of body weights and spleen weights with age-, sex-matched wild type mice revealed significantly lower body weight and pronounced splenomegaly in the ECKO mice at around 2 months of age (Fig. 1A). H&E stained ECKO spleens showed disrupted architecture containing significantly enlarged red pulps and underdeveloped white pulps (Fig. 1B), indicating extensive extramedullary hematopoiesis. Because the extramedullary hematopoiesis is likely to be a compensatory mechanism due to bone marrow dysfunction, Giemsa staining was performed to examine the bone marrows of wild type and ECKO mice. ECKO mice exhibited decreased erythropoiesis with a relatively increased ratio of myeloid to erythroid lineages (Fig. 1C). Consistently, the hematocrit percentage was markedly decreased in ECKO mice compared with wild type mice at 2 months of age (Fig. 1D). This difference was also reflected in the peripheral blood smear showing markedly reduced numbers of red blood cells in ECKO mice compared with wild type controls (Fig. 1E).

**Figure 1.**
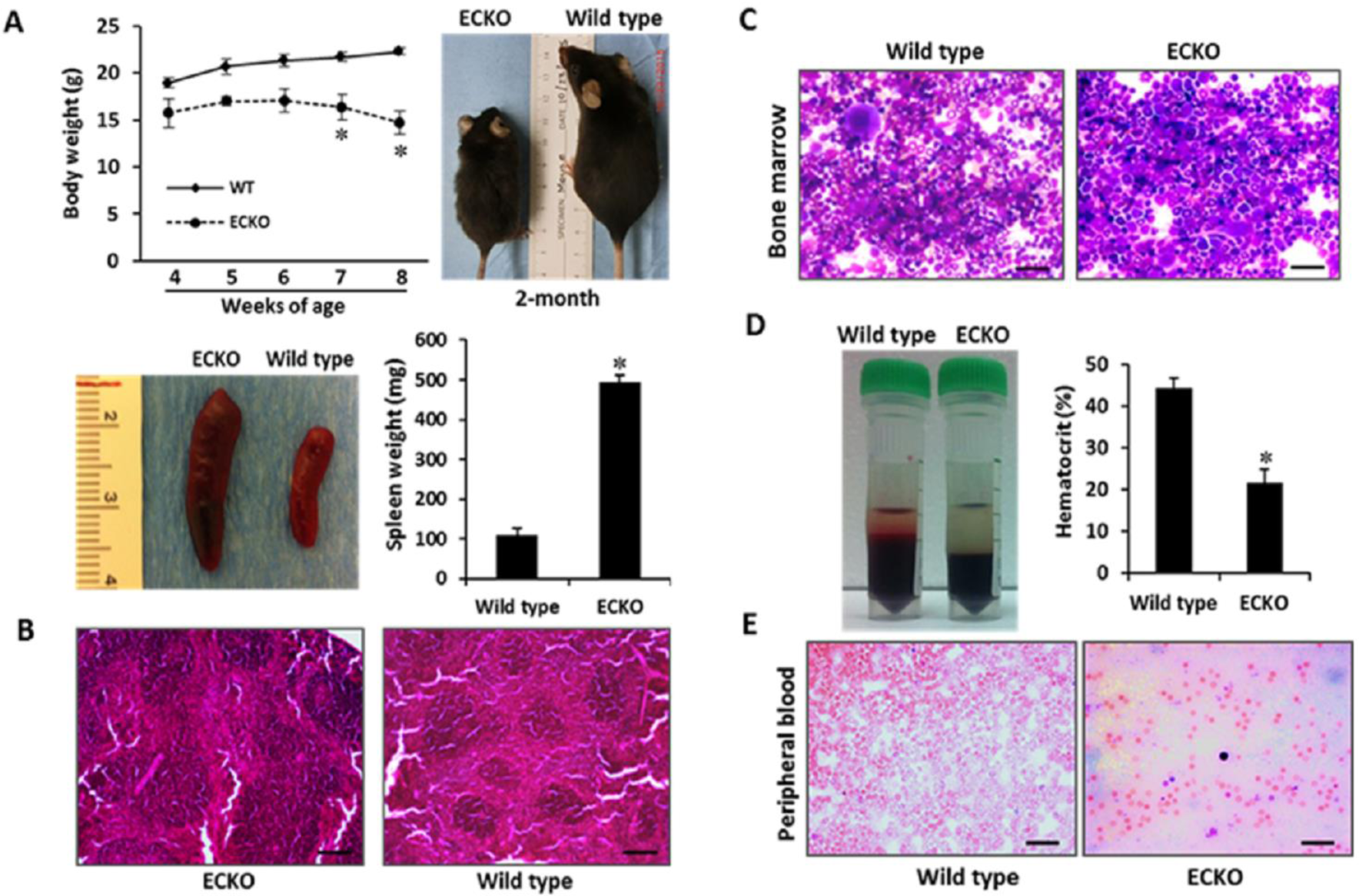
Characterization of mice with EC-specific deletion of MCPIP. (A) Comparison of body weight and spleen weight with age-, sex-matched wild type mice, ECKO mice showed significantly lower body weight and pronounced splenomegaly at around 2 months of age; *\*p* < 0.05, n = 6. (B) H&E stained ECKO spleens showed disrupted architecture containing significantly enlarged red pulps and underdeveloped white pulps. (C) Bone marrow morphology assessed by Giemsa staining showed decreased erythropoiesis with relatively increased ratio of myeloid to erythroid lineages in ECKO mice. (D, E) The percentage of hematocrit and numbers of red blood cells in the peripheral blood smear in ECKO mice was markedly decreased compared with wild type mice at 2 months of age. *\*p* < 0.05, n = 6.

### Serum profiling reveals increased inflammatory response and endothelial dysfunction in the ECKO mice

We performed multiple analytic profiling on serum from wild type and ECKO mice at 2 months of age, when showing splenomegaly, to investigate the impact of MCPIP deletion on endothelial function. Circulating levels of cytokines, chemokines, and biomarkers of endothelial dysfunction, such as sP-selectin and PAI-1 were measured. Of the 32 serum proteins tested, 28 were detectable in both the ECKO mice and age-, sex-matched wild type controls (Table 1). ECHO mice showed elevated levels of IL-2, IL-3, IL-4, IL-5, IL-6, IL-7, IL-12p40, IL-13, IL-15, IL-17, TNF-α, G-CSF, CCL2, CCL11, MIP-1α, MIP-1β, and CXCL9 in comparison with the wild type controls (Table 1). Of the other analytes tested, vascular endothelial growth factor (VEGF), KC/CXCL1, LIF and IP-10 were elevated while LIX/CXCL5 was decreased in the ECKO mice in comparison with the wild type controls (Table 1). Increased levels of soluble adhesion molecules such as sP-selectin and PAI-1, biomarkers of endothelial dysfunction, were significantly elevated in the ECKO mice (Table 1).

**Table 1.** Serum cytokine profiles in the wild type and ECKO mice

|  | Wild type (n = 3) | ECKO (n = 3) | *P* value |
| --- | --- | --- | --- |
| Cytokines (pg/ml) |  |  |  |
| G-CSF | 228.7 ± 11.8 | 707.1 ± 8.2 | < 0.0001 |
| M-CSF | 20.6 ± 9.8 | 59.1 ± 35.7 | 0.146 |
| IL-1β | 19.5 ± 7.1 | 30.2 ± 8.2 | 0.1627 |
| IL-2 | 19.2 ± 5.9 | 221.7 ± 18.8 | < 0.0001 |
| IL-3 | 6.6 ± 2.8 | 18.5 ± 0.8 | 0.0021 |
| IL-4 | 0.2 ± 0.03 | 3.8 ± 0.05 | < 0.0001 |
| IL-5 | 5.8 ± 0.8 | 381.9 ± 8.4 | < 0.0001 |
| IL-6 | 1.0 ± 0.7 | 22.8 ± 0.7 | < 0.0001 |
| IL-7 | 8.1 ± 3.3 | 14.9 ± 1.1 | 0.0276 |
| IL-10 | 6.6 ± 3.1 | 54.8 ± 16.3 | 0.0073 |
| IL-12 p40 | 29.8 ± 5.3 | 71.6 ± 10.7 | 0.0037 |
| IL-13 | 62.6 ± 22.1 | 760.2 ± 102.1 | 0.0003 |
| IL-15 | 97.1 ± 67.7 | 127.6 ± 33.2 | 0.5222 |
| IL-17 | 2.5 ± 0.4 | 6.0 ± 0.3 | 0.0003 |
| TNF-α | 16.1 ± 1.4 | 24.9 ± 5.2 | 0.0473 |
| Chemokines (pg/ml) |  |  |  |
| CCL11 | 482.8 ± 72.1 | 1420.2 ± 126.2 | 0.0004 |
| CCL2/MCP-1 | 59.9 ± 6.7 | 96.6 ± 3.4 | 0.001 |
| MIP-1α | 46.7 ± 38.2 | 335.3 ± 39.2 | 0.0008 |
| MIP-1β | 11.4 ± 4.1 | 244.3 ± 40.5 | 0.0006 |
| CXCL9/MIG | 74.4 ± 8.6 | 142.9 ± 0.9 | 0.0002 |
| RANTES | 43.5 ± 8.1 | 76.4 ± 28.1 | 0.1232 |
| Others (pg/ml) |  |  |  |
| KC | 83.6 ± 6.5 | 524.5 ± 23.5 | <0.0001 |
| LIX | 7302.1 ± 858.9 | 120 ± 45.3 | 0.0001 |
| IP-10 | 59.5 ± 6.7 | 3720.3 ± 212.3 | < 0.0001 |
| LIF | 1.4 ± 0.9 | 3.4 ± 0.3 | 0.0217 |
| VEGF | 1.0 ± 0.4 | 6.9 ± 0.4 | < 0.0001 |
| PAI-1 | 4.6 ± 0.6 | 19.7 ± 0.8 | < 0.0001 |
| sP-selectin | 166.8 ± 4.6 | 226.6 ± 6.9 | 0.0002 |

Results were expressed as mean ± SD. *P* values were shown for comparison with corresponding wild type. G-CSF, Granulocyte-colony stimulating factor; M-CSF, Macrophage colony-stimulating factor; IL, interleukin; TNF-α, Tumor necrosis factor-alpha; CCL, C-C motif chemokine ligand; MIP, Macrophage inflammatory proteins; CXCL, C-X-C motif ligand; RANTES, Regulated on activation, normal T Cell expressed and secreted; KC, Keratinocyte chemoattractant; LIX, C-X-C motif ligand 5; IP-10, Interferon gamma-induced protein 10; LIF, Leukemia inhibitory factor; VEGF, Vascular endothelial growth factor; PAI-1, Plasminogen activator inhibitor-1; sP-selectin, Soluble platelet selectin.

### Endothelial MCPIP deletion causes enhanced vascular permeability and multi-organ inflammation

To test for the impact of endothelial MCPIP deletion on vascular leakage, Evans blue dye was injected in wild type controls and ECKO mice via the tail vein, and various organs (ear, heart, and kidney) were perfused 30 minutes later with saline to clear the remaining dye from the circulation. Ear, heart, and kidney tissues from the ECKO mice exhibited higher levels of dye retention than wild type controls, indicating increased vessel permeability (Fig. 2A). ECKO mice also exhibited an increased ratio of heart-to-body weight compared with the wild type controls (Fig. 2B). H&E-stained heart sections showed marked vascular thrombosis (*arrows*) with cellular infiltration in the myocardium of the ECKO mice (Figs. 2C, D).

**Figure 2.**
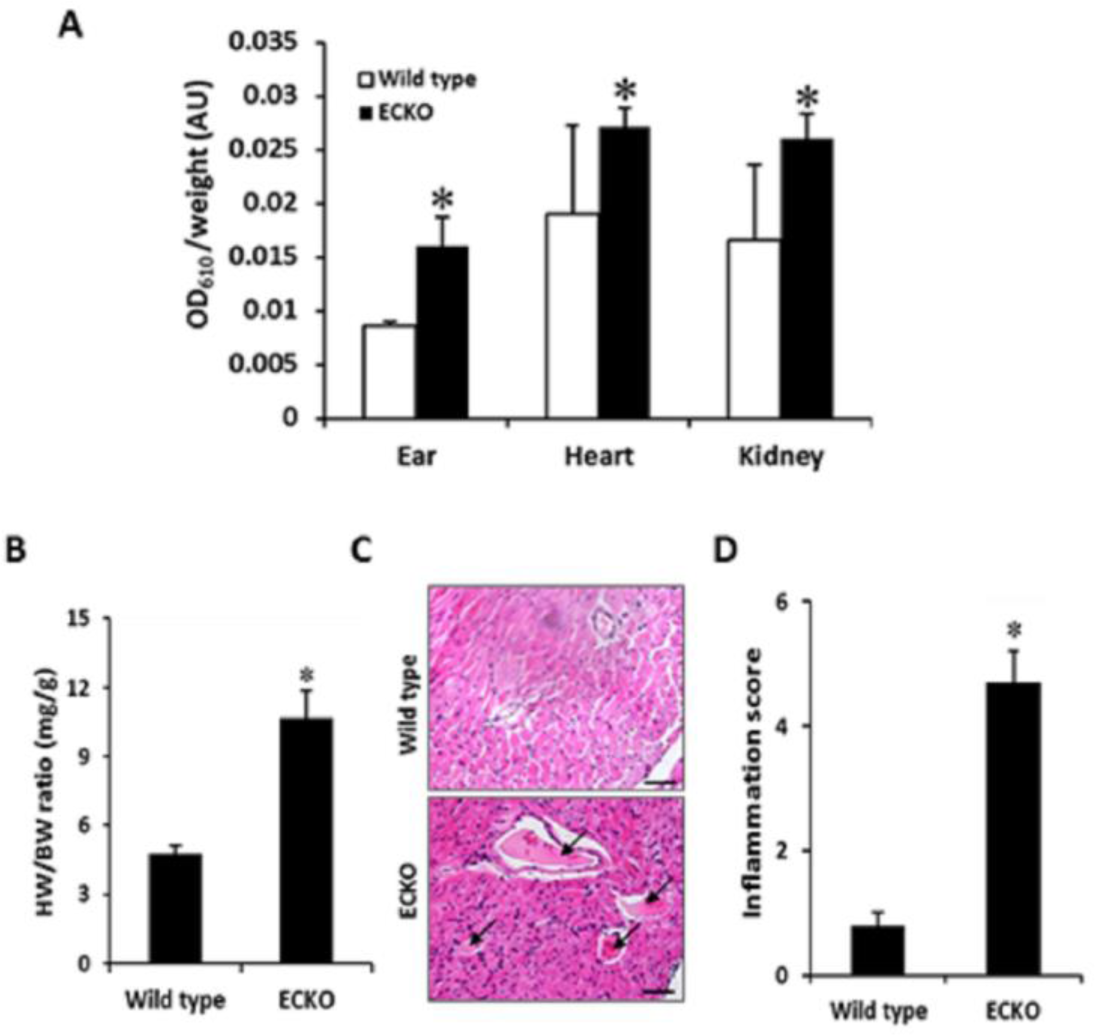
Figure 2. Endothelial-specific MCPIP deletion enhances vascular permeability and cellular infiltration in the myocardium. (A) Vascular permeability assay in vivo by injection of Evans blue dye showed increased dye retention in the ear, heart, and kidney tissues from ECKO mice, indicating increased vessel permeability. *\*p* < 0.05, n = 6. (B) ECKO mice exhibited increased ratio of heart weight (HW) to body weight (BW). *\*p* < 0.05, n = 6. (C, D) H&E-stained heart sections showed marked vascular thrombosis (arrows) with extensive inflammatory cell infiltration in ECKO mice. *\*p* < 0.05, n = 6.

The increased red blood cell extravasation, plasma protein leakage, cellular infiltration, and thrombus with luminal obstruction of blood vessels were also observed in lung tissues of the ECKO mice (Fig. 3A). Cellular infiltrates in both perivascular and peribronchiolar lung tissue were markedly increased in the ECKO mice in comparison to the wild type controls (Figs. 3B, C), which was reflected by the increased ratio of lung weight to body weight and the ratio of lungs’ wet weight to dry weight (Figs. 3D, E). Furthermore, perivascular fibrosis was dominant in the muscular wall of the pulmonary artery which manifested as severe, diffuse intimal hyperplasia with fibrinoid material within the blood vessels of the ECKO mice (Fig. 3A, blue staining). We analyzed the levels of IL-6, a major cytokine that has a diverse role in driving chronic inflammation, EC dysfunction, and fibrogenesis in the lung tissues.20 IL-6 was elevated at 3-week-old ECKO compared with wild type control mice, and became more pronounced in 2-month-old mice (Fig. 3F). Similarly, the levels of adhesion molecules such as ICAM-1, VCAM-1, E-selectin and PAI-1, markers of EC dysfunction, were markedly increased in 2-month-old ECKO mice compared with the wild type controls (Fig. 3G). Consistent with these findings, the ECKO mice subjected to LPS showed significantly higher mortality, with the majority of death occurring in the first 24 hours after LPS challenge (Fig. 3H).

**Figure 3.**
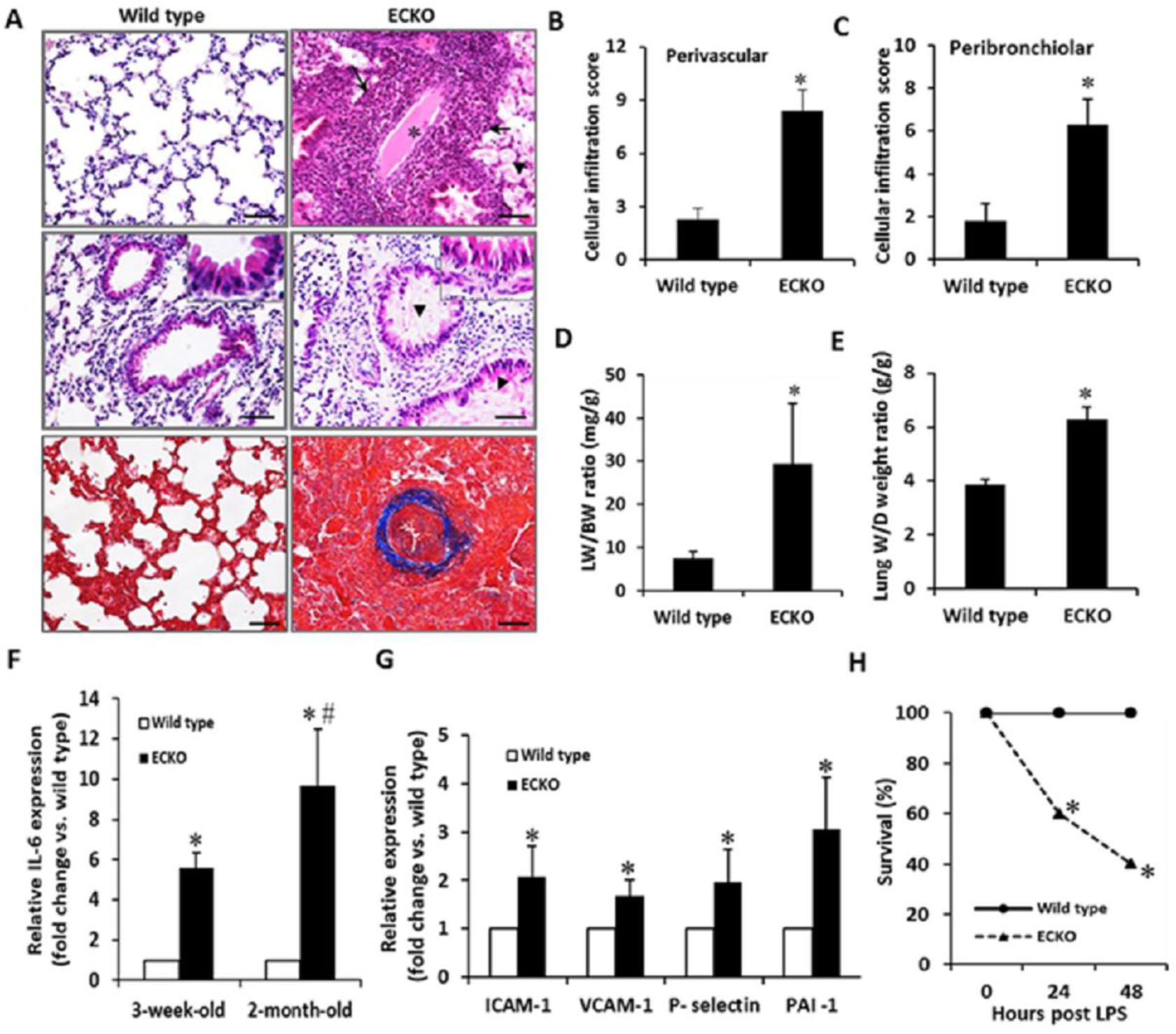
Endothelial-specific MCPIP deletion causes a severe inflammatory response in the lung. (A) H&E-stained lung sections showed increased plasma protein leakage (arrow head), cellular infiltration (arrow), thrombus with luminal obstruction of blood vessels (star), perivascular fibrosis (blue staining) and diffuse intimal hyperplasia in muscular pulmonary artery in lung tissues of the ECKO mice. (B, C) Both perivascular and peribronchiolar lymphocytic inflammatory infiltrates were markedly increased in the lung of ECKO mice. *\*p* < 0.05, n = 6. (D, E) ECKO mice showed the increased ratio of lung wet weight to body weight (LW/BW) and the ratio of lung wet weight to dry weight. *\*p* < 0.05, n = 6. (F) qRT-PCR analysis of IL-6 expression in the lungs at 3 weeks and 2months of age. *\*p* < 0.05, *#p* < 0.01, n = 6. (G) qRT-PCR analysis of expression of ICAM-1, VCAM-1, P-selectin and PAI-1in lungs at 2 months of age. *\*p* < 0.05, n = 6. (H) Survival analysis showed significantly increased mortality of ECKO mice after LPS challenge (25mg/kg). *\*p* < 0.05, n = 10.

Muscle wasting is also an important feature of inflammatory conditions.^21^ The ECKO mice exhibited a profound muscle wasting with age when compared to the age-matched wild type controls. The ECKO mice showed a 60% reduction in muscle fiber cross-sectional area and a 40% reduction in muscle fiber width at 2 months of age (Figs. 4A-C). Histopathological analysis demonstrated markedly increased cellular infiltration in the skeletal muscles of ECKO mice in comparison with the wild type controls (Fig. 4D). This observation was reflected by the increased expression of atrophy-related genes encoding two muscle-specific ubiquitin ligases, Atrogin-1 and MuRF1 (Figs. 4E, F).^22^

**Figure 4.**
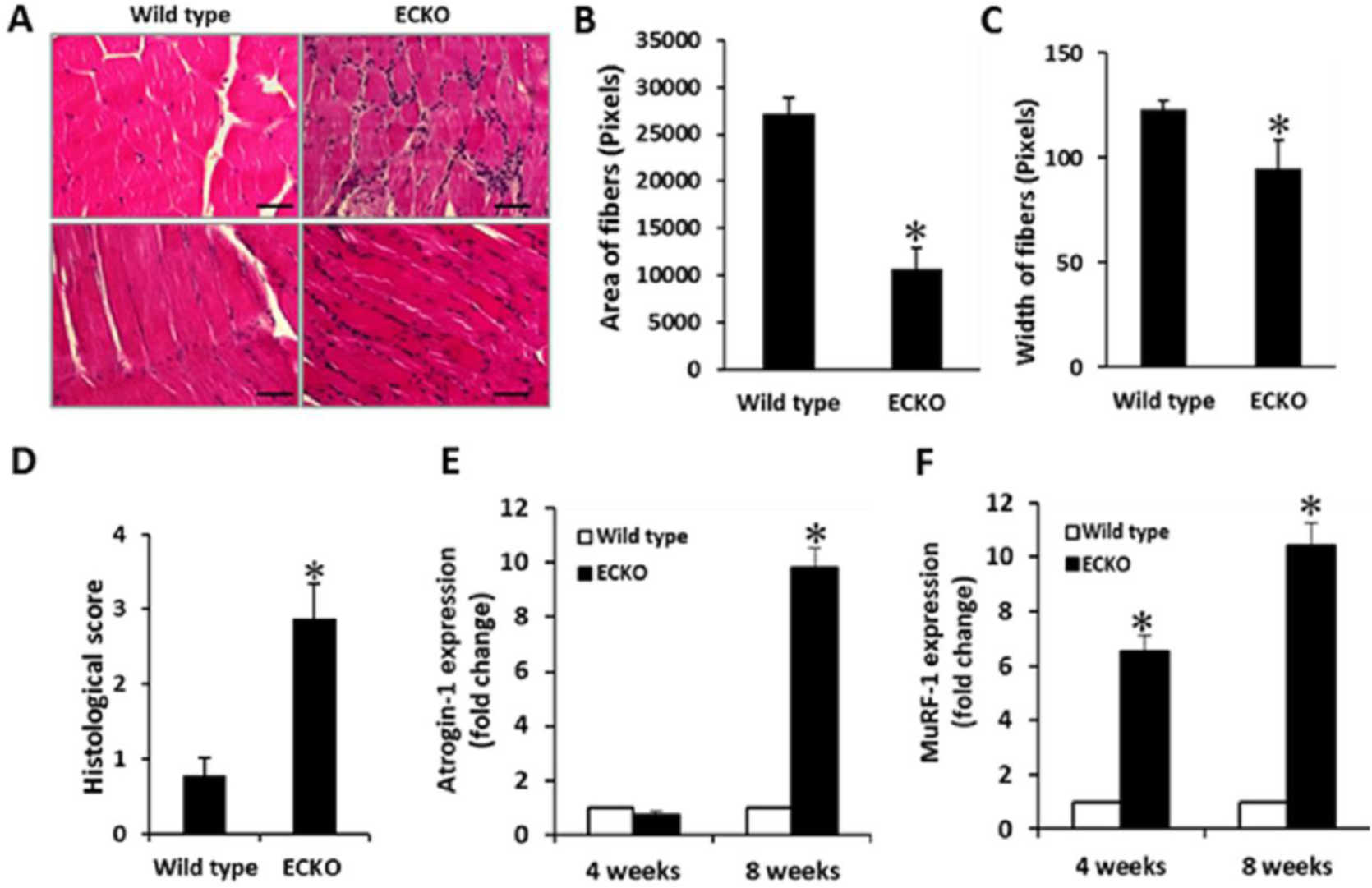
Endothelial-specific MCPIP deletion causes inflammatory muscle wasting. (A) H&E-stained gastrocnemius muscles sections show cellular infiltration and small size muscle fiber in the ECKO mice at 2 months of age. (B, C) Bar graphs show a 60% reduction in muscle fiber cross-sectional area and a 40% reduction in muscle fiber width in ECKO mice at 2 months of age. *\*p* < 0.05, n = 6. (D) A histogram shows cellular infiltration score. *\*p* < 0.05, n = 6. (E, F) qRT-PCR analysis of expression of atrophy-related genes, Atrogin-1 and MuRF1, in the gastrocnemius muscles at 4, and 8 weeks of age. *\*p* < 0.05, n = 6.

### Mice with endothelial MCPIP deletion exhibit reduced hindlimb perfusion and impaired recovery of perfusion post ischemia

To assess the impact of EC-specific MCPIP deletion on vascular function in adult tissues in vivo, we used laser Doppler imaging to assess blood perfusion of the hindlimbs of the 2-month-old ECKO and wild type mice. A significant decrease in blood flow was observed in the ECKO mice in comparison with the wild type controls (Fig. 5A). We then analyzed blood flow recovery in the ECKO and wild type mice using a hindlimb ischemia model. In this model, ischemia was induced by ligation and excision of a segment of the femoral artery. After ligation of the femoral artery, an immediate and marked but equal reduction in perfusion in the ischemic hindlimb was observed in both the wild type and ECKO mice (Figs. 5B, C). Blood flow recovery in the ischemic hindlimb increased over subsequent days in both the wild type and ECKO mice; however, blood flow recovery was significantly impaired in the ECKO mice, which became more apparent 21 days after ligation (Fig. 5C). Consistent with the perfusion data, limb use and appearance scores were significantly worse in the ECKO mice compared with the wild type controls (Figs 5D, E).

**Figure 5.**
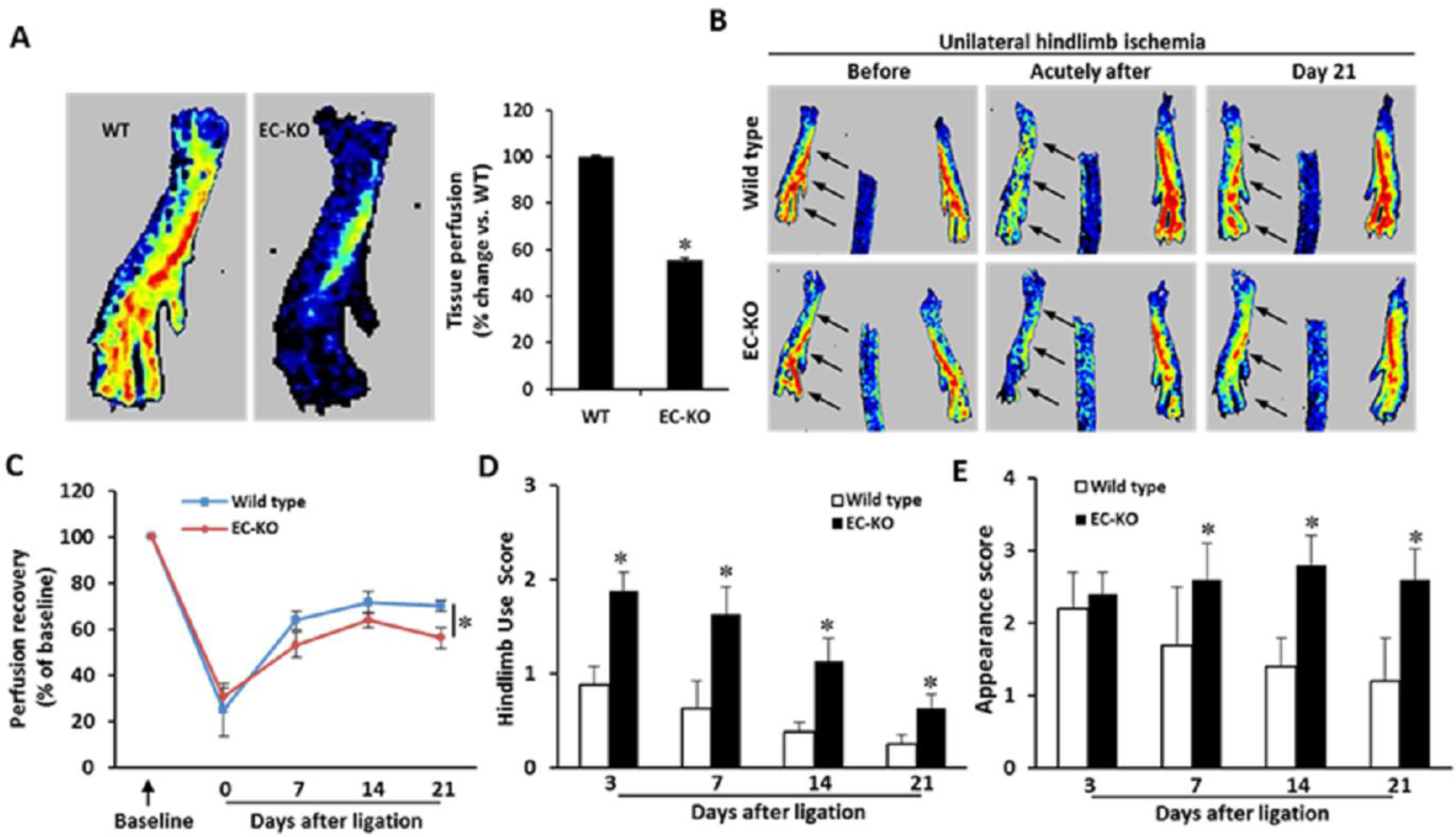
ECKO mice have impaired perfusion after limb ischemia. (A) Representative laser Doppler images of plantar foots of wild type and ECKO mice show a significant decline in blood perfusion in ECKO mice. *\*p* < 0.05, n = 6. (B, C) Wild type and ECKO mice were subjected to hindlimb ischemia. Blood flow recovery was measured by relative values of foot perfusion between ischemic and non-ischemic legs. Representative laser Doppler perfusion images of plantar foot show decreased perfusion recovery after ligation in the ECKO mice. *002A;p* < 0.05, n = 10. (D, E) ECKO mice have greater use score and appearance score. *\*p* < 0.05, n = 10.

### Postnatal angiogenesis is impaired in endothelial-specific MCPIP knockout mice

Lower blood perfusion in the ECKO mice could indicate reduced postnatal angiogenesis. Therefore, we examined lumen diameter and wall thickness of collaterals to assess the impact of endothelial-specific MCPIP deletion on angiogenesis by histomorphometric analysis at baseline (non-ligated) and at day 21 after ligation (ligated). ECKO mice exhibited reduced lumen diameters with thrombus in blood vessels, whereas no thrombus with no luminal obstruction of blood vessels was found in wild type mice (Figs. 6A-C). 21 days after ligation, wild type mice displayed increased collateral diameter and wall area in the ischemic tissues; however, this enhancement was markedly attenuated in the ECKO mice (Figs. 6A-C). Additionally, capillary density was measured with anti-CD31 antibody in non-ligated and ligated limb tissues with immunofluorescence. ECKO mice had an increased, morphologically disorganized capillary distribution in comparison with the wild type controls (Figs. 6D, E). These data, as well as the reduced blood flow before ligation and impaired recovery of perfusion after ligation, suggest that vasculature in ECKO mice has an impaired ability to remodel into fully functional vessels due to MCPIP deficiency in endothelial cells. This hypothesis is supported by histological analysis revealing significantly increased areas of injury and necrosis after femoral artery ligation in the ECKO mice when compared with the wild type controls (Figs. 6F, G).

**Figure 6.**
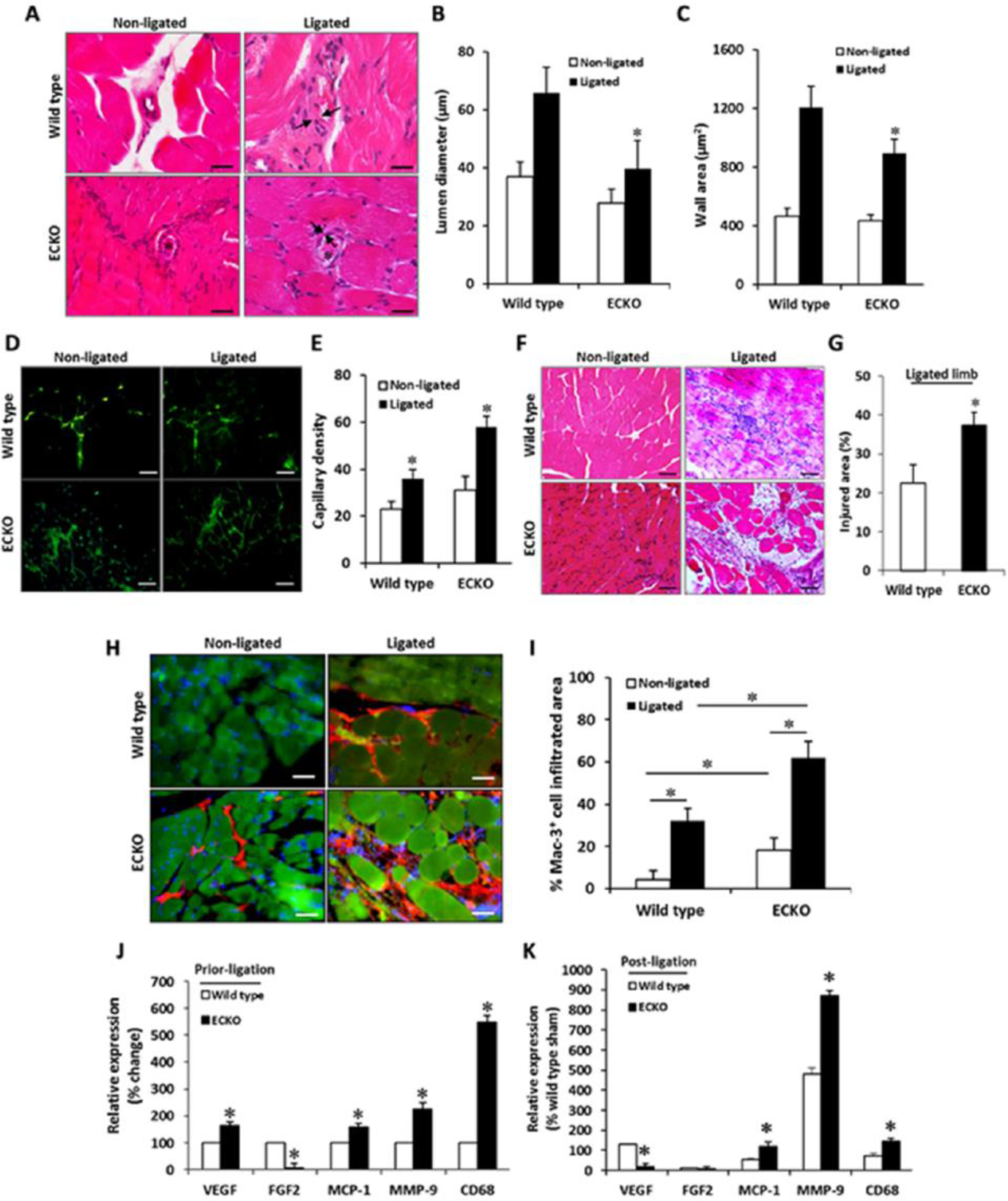
Endothelial-specific MCPIP deletion impairs post-ischemic neovascularization in ECKO mice. (A) Representative H&E-stained sections show collateral remodeling after hindlimb ischemia between wild type and ECKO mice. Arrows indicate collateral wall. Stars indicate thrombosis within collateral lumen. (B, C) The luminal diameter and wall area calculated from the measurements of luminal and perivascular tracing shows impaired collateral growth in the ECKO mice. *\*p* < 0.05, n = 6. (D, E) Ischemic muscles at day 21 were immunostained with endothelial cell marker CD31 (green), which shows increased capillary density in ECKO mice. *\*p* < 0.05, n = 6. (F, G) Representative H&E-stained sections of ischemic muscles show the increased percentage of injury area with cellular infiltration in the ECKO mice. *\*p* < 0.05, n = 6. (H, I) The cellular infiltration in the ischemic muscles were analyzed by immunostaining with a macrophage marker Mac-3 (red) and quantified as the percentage of Mac-3 positive infiltrated area in the graph. *\*p* < 0.05, n = 6. (J, K) qRT-PCR analysis for expression of VEGF, FGF2, MCP-1, MMP-9, and CD68 in the muscles prior to ischemia and post-ischemic 21 days. *\*p* < 0.05, n = 6.

### Endothelial MCPIP deletion affects expression of cytokines and angiogenic factors

To gain a better understanding of how endothelial-specific MCPIP deletion influences postnatal angiogenesis, we examined cellular infiltration in non-ligated and ligated legs of wild type and ECKO mice by immunofluorescence against Mac-3, a marker of macrophages. Basic accumulation of Mac-3 positive macrophages (red fluorescence) in the non-ligated limb was more extensive in the ECKO mice compared with the wild type controls (Fig. 6H). At 21 days post ligation, accumulation of Mac-3 positive macrophages in the ischemic muscles increased 3-fold in the ECKO mice in comparison with the wild type controls (Fig. 6I). Expression of CD68 and MMP-9 in both non-ligated and ligated tissues significantly increased in the ECKO mice compared to the wild type controls (Figs. 6J, K). Accumulation of macrophages is driven mainly by chemokine MCP-1.^23^ The level of mRNA for MCP-1 in both non-ligated and ligated muscles showed significantly higher levels in the ECKO mice compared with the wild type controls (Figs. 6J, K).

MCPIP has been shown to regulate expression of multiple genes encoding angiogenic factors such as VEGF and FGF-2.^19^ We examined whether endothelial MCPIP deletion-induced vascular phenotype in ECKO mice could be explained, at least in part, by the altered expression of these genes. Expression for FGF-2 in both non-ischemic and ischemic muscles was greatly reduced in the ECKO mice compared to the wild type controls (Figs. 6J, K). The level of mRNA for VEGF in ischemic tissues was also significantly lower in ECKO mice, although its level in non-ischemic tissues was higher in ECKO mice than in the wild type controls (Figs. 6J, K).

### ECKO mice display impaired vessel sprouting and wound healing

Given the importance of systemic factors such as blood pressure, blood flow, and inflammatory reactions on postnatal angiogenesis, we performed a mouse aortic ring assay ex vivo using isolated aortas from the ECKO mice or their corresponding wild type controls to determine the impact of endothelial MCPIP deletion in capillary sprouting from pre-existing vessels. Mouse thoracic aortas were sectioned, embedded in Matrigel, and cultured with EBM medium, and neovessel sprouts were blindly counted on day 6. In contrast to the wild type aortic rings from which microvessels sprouted, aortas from the ECKO mice displayed markedly impaired capillary sprouting and tube elongation (Figs. 7A, B), indicating MCPIP deletion in endothelial cells impairs angiogenesis.

**Figure 7.**
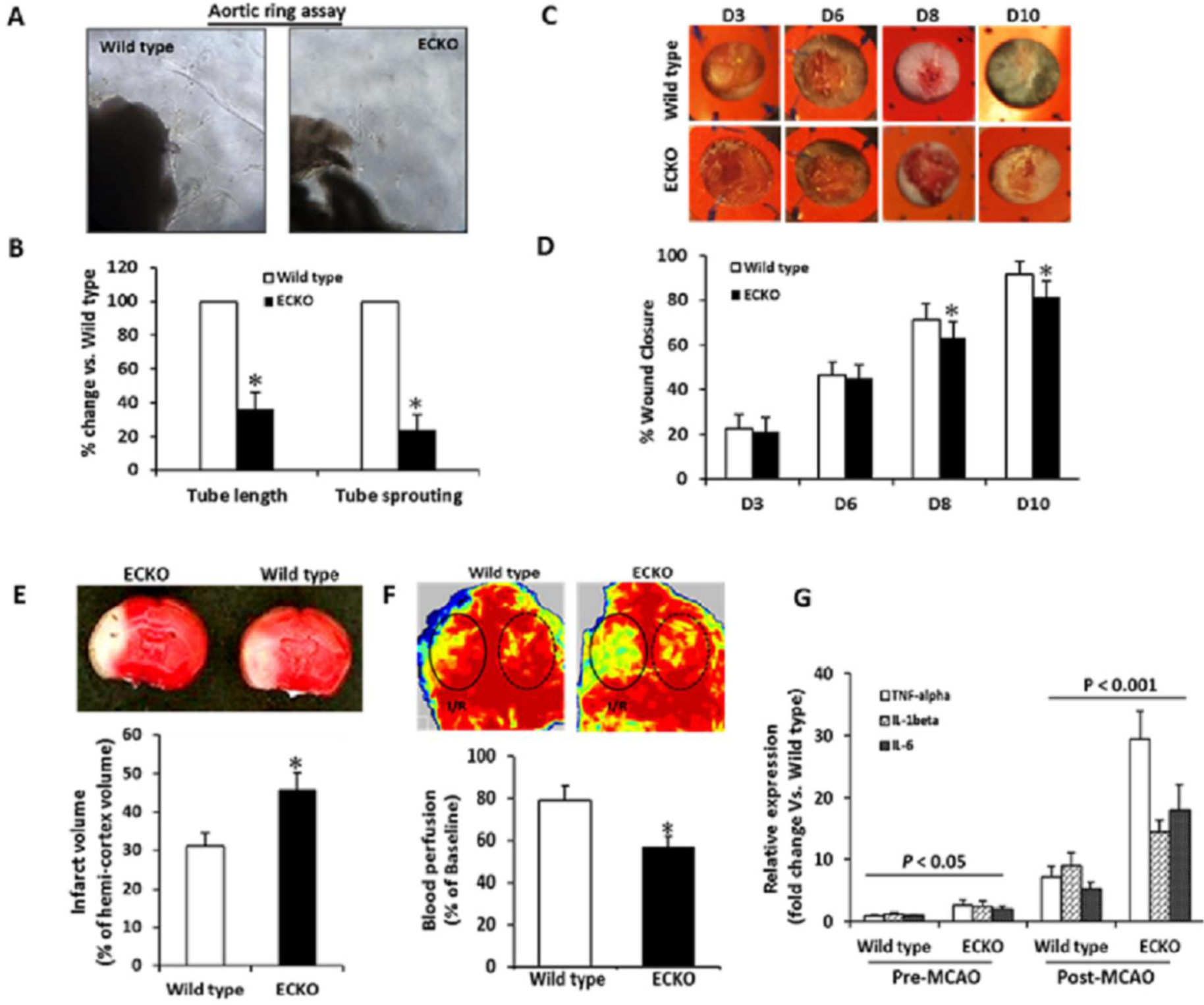
ECKO mice have impaired vessel sprouting, wound healing, and larger cerebral infarction after MCA occlusion. (A, B) Aortas were harvested from 2-month-old ECKO and their littermate controls and cultured in Matrigel for 6 days. Capillary sprouting and tube length were quantified in 6 aortic rings under microscopy. *\*p* < 0.05, n = 6. (C, D) Representative images of wound healing in ECKO and their littermate controls and imaged at the indicated times, and quantification of the area of wound closure is shown in bar graph. *\*p* < 0.05, n = 6. (E) Coronal slices after 2h of MCA occlusion followed by 24h of reperfusion show infarcted area (Upper panel, pale region), and quantification of the infarct volume is shown in bar graph (Lower panel). *\*p* < 0.05, n = 5. (F) Laser Doppler perfusion images show impaired recovery of perfusion (Upper panel, circled region) after 2h of MCA occlusion followed by 24h of reperfusion, and quantification of perfusion recovery is shown in the bar graph (Lower panel). *\*p* < 0.05, n = 5. (G) qRT-PCR analysis for expression of TNF-α, IL-1β, and IL-6 in the brain tissues harvested from 2-month-old ECKO and their littermate controls prior to MCA and post- ischemia/reperfusion. n = 5.

Since the rate of wound healing reflects the underlying angiogenesis, ^24^ we subsequently examined the effect of EC-specific MCPIP deletion on wound healing using a puncture wound healing model. Full-thickness excisional wounds (6 mm) were made on the dorsum of anesthetized ECKO mice or on their corresponding wild type mice. Visual inspection and quantitative analysis demonstrated that wound healing in ECKO mice was markedly delayed compared with the wild type controls at the same time points after day 7 (Figs. 7C, D).

Because infarct volume upon middle cerebral artery (MCA) occlusion is known to correlate with pial collateral density in mice, ^25^ we performed MCA occlusion for 2 hours followed by reperfusion for 24 hours, and measured infarct volume in the ECKO mice and their wild type littermates. ECKO mice had about 1.5-fold more infarct volume compared with their wild type littermates (Fig. 7E), indicating that the formation of native collaterals is impaired in ECKO mice. This was reflected by the decreased blood perfusion to the brain as assessed by laser Doppler perfusion imaging (Fig. 7F). Moreover, increased levels of proinflammatory cytokines such as TNF-α, IL-1, and IL-6 were observed in the brain tissues of ECKO mice at both pre- and post-occlusion of the MCA as compared with their wild type littermates (Fig. 7G).

### MCPIP deficiency induces NF-κB activity and pro-inflammatory miRNAs in ECs in vitro and in vivo

NF-κB activation is critical to inflammatory gene expression in endothelial cells, ^4^ and MCPIP was reported to regulate inflammatory signaling by suppressing NF-κB activation.^11^ To test whether the constitutive level of MCPIP present in ECs would suppress NF-κB activation and thus keep ECs in a quiescent state, we utilized siRNA knock down in HUVECs to assess the potential function of the constitutive level of MCPIP in ECs. Treatment with MCPIP specific siRNA elevated the levels of NF-κB activity by 4.6-fold (Fig. 8A). This activation was also reflected by the increased phosphorylation of IκB in the lung tissue of ECKO mice (Fig. 8B). As expected from NF-κB activation, the mRNA levels of NF-κB-dependent cytokines (e.g., IL-1, IL-6, MCP- 1, TNF-α) and pro-coagulants (e.g., PAI-1, TF) were higher in HUVECs transfected with MCPIP specific siRNA than in cells transfected with the scramble siRNA (Figs. 8C, D). To verify the observed effects of MCPIP in vivo on the expression of adhesion molecules and pro-coagulants, heart and lung tissues from adult ECKO mice and their littermates were examined for expression of ICAM-1, VCAM-1, P-selectin, PAI-1 and TF by real-time quantitative PCR. The mRNA levels of ICAM-1, VCAM-1, P-selectin, PAI-1, and TF in the lung and heart tissue were markedly increased in the ECKO mice compared with their wild type littermates (Figs. 8G, E). The levels of mRNA for inflammatory cytokines was also significantly elevated in the myocardium of ECKO mice when compared to their wild type littermates (Fig. 8F).

**Figure 8.**
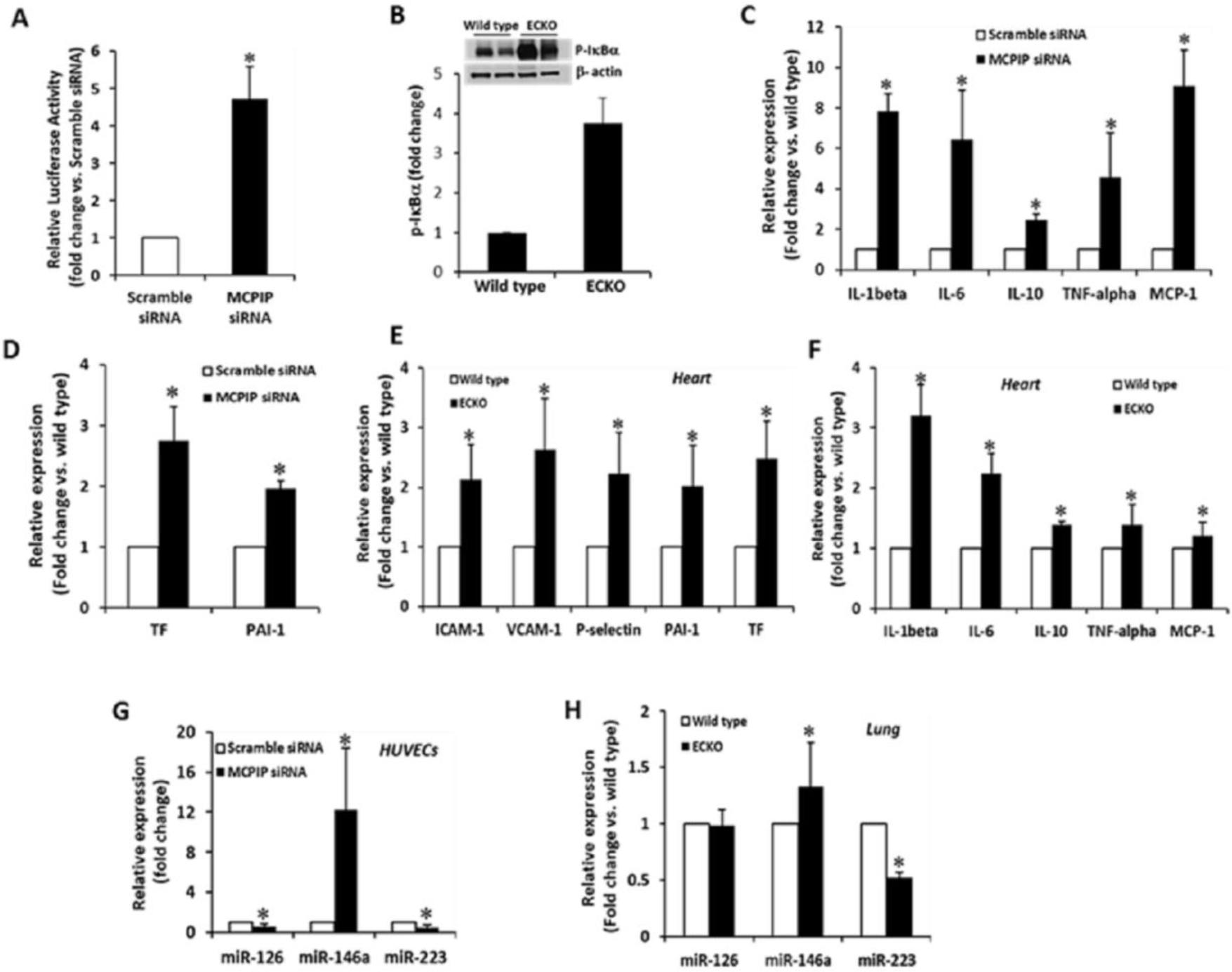
Endogenous MCPIP restrains endothelial activation in vitro and in vivo. (A) HUVECs were transfected with a MCPIP specific siRNA or scramble siRNA for 24h, and the levels of NF-κB activity were assayed by measurement of NF-κB promoter-luciferase reporter activity. *\*p* < 0.05. (B) Activation of NF-κB in vivo was assessed by measuring the levels of phosphorylated IκBα in the lungs of ECKO mice and their littermate controls by immunoblots. β-actin was used as a loading control. *\*p* < 0.05. (C, D) The expression of inflammatory genes (IL-1β, IL-6, IL-10, TNF-α, and MCP-1) and pro-coagulants (TF and PAI-1) in MCPIP-specific siRNA and scramble siRNA-treated HUVECs were assessed by qRT-PCR. *\*p* < 0.05. (E, F) Levels of adhesive molecules (ICAM-1, VCAM-1, and P-selectin), pro-coagulants (TF and PAI-1), and cytokines (IL-1, IL-6, IL-10, TNF-α, and MCP-1) in hearts of the ECKO mice and their littermate controls were assessed by qRT-PCR. *\*p* < 0.05. (G) Knockdown of MCPIP inhibited the expression of miR-126, −223, and increased expression of miR-146a in HUVECs, assayed by qRT-PCR. *\*p* < 0.05. (H) qRT-PCR analysis of miR-126, −223, and −146a expression in lungs of 2-month-old ECKO mice and their littermate controls. *\*p* < 0.05. Experiments were repeated three times.

Several microRNAs such as miR-126, −146a, and −223 were previously reported to be highly enriched in ECs, and their dysregulation has been shown to be involved in EC dysfunction.^6-8^ We assessed the expression of these microRNAs in HUVECs in response to knock down of MCPIP, and found that the levels of miR-146a were markedly elevated and that miR-126 and −223 were reduced by knock down of MCPIP (Fig. 8G), suggesting expression of these microRNAs in HUVECs is regulated by the constitutive level of MCPIP. Next, we assessed the expression of these microRNAs in the lung tissue of 2-month-old ECKO mice and age-, sex-matched wild type controls. The results showed elevated levels of miR-146a, reduced levels of miR-223, and no change in the levels of miR-126 in the lung tissues of ECKO mice when compared with their wild type littermates (Fig. 8H). These results suggest that the constitutive level of MCPIP in ECs is able to keep ECs in a quiescent state by negatively regulating NF-κB activation and microRNA synthesis.

## Discussion

Molecular mechanisms that control EC quiescence remain largely unknown. In this study, we show for the first time that endothelial MCPIP deficiency causes systemic inflammation, vascular leakage, cellular infiltration, microvascular thrombosis, an enlarged spleen, anemia, muscle wasting and postnatal lethality in mice. These mice also manifested impaired angiogenesis, blood perfusion, and tissue repair after ischemia. Deletion of MCPIP in ECs caused NF-κB-dependent pro-inflammatory signaling and dysregulation of synthesis of microRNAs (e.g., miR-146a, −223) in ECs in vitro and in the lung tissue of ECKO mice in vivo. This is the first report providing direct genetic evidence for a critical role of MCPIP present in ECs in maintaining EC function and in restraining vascular inflammation.

An activated endothelium produces various factors, including proinflammatory cytokines, chemokines, vascular endothelial growth factor (VEGF), and adhesion molecules such as intracellular adhesion molecule-1 (ICAM-1) and P-selectin.^3,4^ In addition, activated ECs can function as antigen-presenting cells by expressing the major histocompatibility complex class II and costimulatory molecule, which induce migration of lymphocytes across endothelium to underlying tissue at the site of vascular damage.^1^ Our findings that loss of MCPIP in ECs in vivo leads to induction of adhesion molecules (e.g., ICAM-1, VCAM-1, P-selectin), chemokines (e.g., CCL11, CCL2), cytokines (e.g., IL-1β, IL-6), and other NF-κB -dependent mediators (e.g., PAI-1, TF) suggest that the constitutive activity of MCPIP serves to suppress NF-κB -mediated EC activation and keep ECs in a quiescent state. In addition, NF-κB activity and expression of NF-κB target genes in the heart and lung were significantly increased in the ECKO mice with increased leukocyte infiltration, excessive vascular leak and thrombosis, and increased sensitivity to LPS as demonstrated by increased mortality resulting from LPS challenge, when compared with their wild type littermates. Supporting this observation is the finding that NF-κB inhibitor MG-132 induced MCPIP expression, and knockdown of MCPIP increased NF-κB activity and expression of NF-κB target genes.^26^ As further support, knock down MCPIP in ECs increased the phosphorylation of eNOS and NO production,^17^ increased expression of VCAM-1 and the adhesion capabilities of THP-1 cells to HUVECs, while overexpression of MCPIP inhibited TNF-α-induced expression of VCAM-1 and THP-1 cell adherence to HUVECs.^16^

Several studies have demonstrated the role of microRNAs in inflammation and immune response in ECs.^6-8^ miRNA-146a, reported to be enriched in ECs, represses EC activation by inhibiting pro-inflammatory signaling of NF-κB.^27^ miRNA-126, an EC-specific microRNA, was reported to regulate EC expression of adhesion molecules like VCAM-1 in HUVECs.^28^ Inhibition of miRNA- 126 decreases tube formation and sprouting of ECs, as well as migration and proliferation in response to VEGF.^29,30^ miRNA-223 was shown to inhibit NF-κB activation in ECs and its downregulation promotes glomerular EC activation.^31^ MCPIP was reported to be able to cleave the terminal loops of precursor microRNAs, thus suppressing the biosynthesis of mature miRNAs.^13^ We found that deletion of MCPIP in ECs in mice resulted in increased levels of miRNA-146a, decreased miR-223 without affecting the expression of miR-126 in the lung. Knock down of MCPIP in HUVECs in vitro showed similar changes in the levels of these three microRNAs. These results suggest that anti-Dicer activity of MCPIP may play a role in keeping ECs in a quiescent state via regulating miRNA synthesis. The transcription of microRNAs is also mediated by activation of NF-κB signaling pathway, while some microRNAs have been demonstrated to alter the transcription of other miRNAs.^6^ We found that deletion of MCPIP in ECs *in vivo* and knock down of MCPIP *in vitro* increased the expression of miRNA-146a, whose transcription is mediated, to a large extent, by NF-κB activation.^32^ Thus, up-regulation of miRNA- 146a in the inflamed ECs initiated by MCPIP deletion is likely to form a feedback loop to restrain endothelial activation. We, like others in the past, also observed that induction of miRNA-146a is suppressed by overexpression of MCPIP in the myocardium, ^33^ and in systemic lupus erythematosus.^34^ miR-223 was reported to inhibit ICAM-1 and TF expression in vascular endothelial cells.^35, 36^ Consistent with these reports is our finding that ECKO mice had decreased expression of miR-223, and increased expression of ICAM-1, VCAM-1 and TF in the heart and lung tissues. Taken together, it is likely that the constitutive level of MCPIP in ECs keep them in a quiescent state by inhibition of NF-κB activation via its deubiquitinase activity and by regulation of microRNA synthesis via its anti-Dicer activities.^11, 13^

Accumulating studies have indicated the involvement of NF-κB signaling pathway and microRNAs in angiogenesis.^37^ Inactivation of Dicer, that generates microRNA from its precursor, has been shown to cause defects in vascular development.^38^ Knockdown of Dicer in ECs in vitro was found to reduce the expression of CXCL1 and upregulate endothelial genes, like eNOS and Tie2, indicating that Dicer promotes inflammatory activation and impairs endothelial differentiation.^39^ We previously demonstrated that small interfering RNA–mediated MCPIP knockdown in HUVECs impairs in vitro proangiogenic properties (proliferation, migration, tube formation) and angiogenesis-associated genes.^19^ Consistent with these findings, the aortic rings from the ECKO mice displayed impaired capillary sprouting and tube elongation. ECKO mice failed to mount an adaptive angiogenic response, leading to impaired recovery of perfusion and increased infarct volume after ischemia. These results strongly imply a critical role of the constitutive activity of MCPIP in ECs for angiogenesis.

Despite the complexity of post-ischemic neovascularization, cellular infiltration is critical for post-ischemic neovascularization and collateral vessel remodeling.^40^ In line with this notion, ECKO mice had increased tissue Mac-3-positive myeloid cells and baseline capillary density (CD30^+^ cells), which is associated with an increase in MCP-1, CD68, MMP-9, and VEGF expression in the skeletal muscles. However, enhanced CD31^+^ cells observed in the tissues of ECKO mice did not constitute functional vessels as indicated by the lack of blood perfusion. This is important to note because these cells cannot transdifferentiate into mature endothelial cells needed for functional neo-vascular formation probably due to lack of MCPIP that is known to be required for EC transdifferentiation.^41^ On the other hand, although VEcadherin5-Cre transgenic mice were originally developed as a tool for EC–specific gene deletion, the VEcadherin5 promoter is active in the bone marrow cells and peripheral blood cells.^42^ Thus, there is possibility that MCPIP deletion may occur in some hematopoietic lineages such as myeloid cells, resulting in impaired monocytic transdifferentiation into endothelial-like cells and angiogenesis. Accordingly, the impaired perfusion, larger infarct volume, impaired post-ischemic flow recovery and wound healing in the ECKO mice reflect the immature capillary networks and diminished arterial blood flow recovery due to lack of MCPIP in these cells.

In conclusion, the present study with EC-specific deletion of MCPIP provides the first in vivo evidence for the role of MCPIP in keeping ECs in a quiescent state. Absence of MCPIP induces NF-κB activation and dysregulated microRNA synthesis, resulting in induction of adhesion molecules, pro-inflammatory mediators, and the pro-coagulants. These changes facilitate the rolling, adhesion, activation, and emigration of circulating leukocytes across the vascular wall, thus causing vascular inflammation and dysfunction, resulting in systemic inflammation, impaired angiogenesis and vascular thrombosis, leading to the phenotypic defects we describe in the ECKO mice. Our work collectively highlights a previously unrecognized role of MCPIP in control of EC homeostasis and function. Our previous work,^33^ coupled with the present study, suggests that therapeutic elevation of MCPIP in endothelium may represent a novel approach to prevent the initiation and progression of chronic inflammatory disorders such as cardiovascular diseases, diabetes mellitus and other inflammatory syndromes.

## Acknowledgments

The authors thank Dr. Candice Sareli, for her careful and critical reading of the manuscript, and Miss Maryann Frost, for her care of the animals used in this study.

